# Multisensory integration-attention trade-off in cochlear-implanted deaf individuals

**DOI:** 10.1101/2020.11.17.384586

**Authors:** Luuk P.H. van de Rijt, A. John van Opstal, Marc M. van Wanrooij

## Abstract

The cochlear implant (CI) allows profoundly deaf individuals to partially recover hearing. Still, due to the coarse acoustic information provided by the implant, CI users have considerable difficulties in recognizing speech, especially in noisy environments. CI users therefore rely heavily on visual cues to augment speech comprehension, more so than normal-hearing individuals. However, it is unknown how attention to one (focused) or both (divided) modalities plays a role in multisensory speech recognition. Here we show that unisensory speech listening and reading were negatively impacted in divided-attention tasks for CI users - but not for normal-hearing individuals. Our psychophysical experiments revealed that, as expected, listening thresholds were consistently better for the normal-hearing, while lipreading thresholds were largely similar for the two groups. Moreover, audiovisual speech recognition for normal-hearing individuals could be described well by probabilistic summation of auditory and visual speech recognition, while CI users were better integrators than expected from statistical facilitation alone. Our results suggest that this benefit in integration comes at a cost. Unisensory speech recognition is degraded for CI users when attention needs to be divided across modalities. We conjecture that CI users exhibit an integration-attention trade-off. They focus solely on a single modality during focused-attention tasks, but need to divide their limited attentional resources in situations with uncertainty about the upcoming stimulus modality. We argue that in order to determine the benefit of a CI for speech comprehension, situational factors need to be discounted by presenting speech in realistic or complex audiovisual environments.

**Significance statement:** Deaf individuals using a cochlear implant require significant amounts of effort to listen in noisy environments due to their impoverished hearing. Lipreading can benefit them and reduce the burden of listening by providing an additional source of information. Here we show that the improved speech recognition for audiovisual stimulation comes at a cost, however, as the cochlear-implant users now need to listen and speech-read simultaneously, paying attention to both modalities. The data suggests that cochlear-implant users run into the limits of their attentional resources, and we argue that they, unlike normal-hearing individuals, always need to consider whether a multisensory benefit outweighs the unisensory cost in everyday environments.

## Introduction

Speech comprehension is a challenging task. First, the speech signal itself might be hard to recognize due to poor pronunciation, semantic ambiguities and highly variable and rapid articulation rates (>200 words/min^1^). Second, in common everyday environments, even highly salient speech signals are frequently embedded in acoustic background noise and are masked by other talkers. During face-to-face conversation, non-acoustic cues from seeing a talker’s mouth can improve speech recognition in those situations, through the integration of visual and auditory information^2–6^.

Multisensory integration is beneficial for normal-hearing and normally sighted individuals, whenever multisensory stimuli are in spatial-temporal congruence. The effects of audiovisual integration include behavioral benefits such as shorter response-reaction times^7–9^, increased accuracy and precision^7,10^, better selection, and reduced ambiguity^11^. At the neuronal level, these effects are typically reflected by enhanced activity^9,12,13^. This also applies to more complex auditory stimuli; supplemental visual input enhances speech perception, and audiovisual speech recognition embedded in noise is considerably better than for auditory speech alone^14–17^. The necessity to integrate non-acoustic information to improve performance becomes especially clear for individuals with hearing impairments, such as profoundly deaf individuals using a cochlear implant (CI). The CI typically recovers hearing to an extent that allows the CI user to understand speech in quiet situations, yet creates significant problems under more challenging listening conditions (e.g., noisy surroundings). In these cases, the CI user should rely more on the information obtained from lip reading. Evidence suggests that CI users are indeed better able to integrate visual information with the perturbed acoustic information than normal-hearing individuals^18,19^.

Due to all the observed benefits of multisensory integration, one may forget that it requires paying attention to multiple sensory modalities at the same time. Attention is a neural mechanism by which the brain is able to effectively select a relevant signal from a multitude of competing sources (e.g., finding someone with a red coat in a busy street). When attention is fully focused on a particular sensory modality, say auditory, performance in auditory selection tasks will markedly increase, but visual stimuli will likely be missed, because attention has limited capacity. The opposite occurs when attention is focused on vision. In natural environments, however, the most relevant sensory modality of a potential target may not be known in advance, and therefore focusing attention on a single sensory modality may not be an optimal strategy to maximize perceptual performance. Instead, in such cases, attention should be divided across the relevant modalities. In case of speech perception, these modalities are auditory (listening) and visual (lipreading) signals. Dividing attention across modalities will allow the brain to integrate the multimodal signals when they originate from the same source, and filter out the perturbing background from unrelated sources.

However, because of its limited capacity, dividing attention in an uncertain sensory environment may lead to decreased performance for stimuli that happen to be unisensory, as each modality will receive less attentional amplification than during a fully focused attention task. Here we compared word-recognition performance during focused and divided attention tasks for CI users and normal-hearing individuals, by presenting unisensory and/or bi-sensory spoken sentences in different sensory-noise regimes. Because CI users have more difficulty to process the perturbed auditory input, more effort (i.e., more attention) will be required to understand auditory speech. Therefore, we reasoned that in a divided-attention task, the lapse in attention to audition (and vision) may lead to poorer unisensory performance scores in CI users. In principle, the same reasoning may hold for normal-hearing participants. So far, it remains unclear from the literature whether CI users can successfully divide their attention across modalities, and whether divided attention affects their speech-comprehension abilities.

## Results

### Overview

Fourteen normal-hearing participants and seven post-lingually deaf unilateral implanted CI users had to identify 50 words (see Methods), presented in 155 unique five-word sentences, by selecting the words they recognized (10-alternative, open-ended choice) on a screen. The speech material has been used in a previous study^20^, in which further details about the material can be found. The stimuli were either presented in two separate unisensory, focused-attention blocks, or in one divided-attention block. We varied task difficulty in both experiments, by blurring the video, and by presenting acoustic background noise at several levels.

In the focused-attention experiment (Fig. 1; purple), the sentences were either presented in an acoustic-only block (Fig. 1A,C, purple circles), or in a visual-only block (Fig. 1B,D, purple circles), in which the participant could focus solely on listening or lipreading, respectively. In the divided-attention experiment auditory (Fig. 1A,C, green diamonds), visual (Fig. 1B,D, green diamonds) and audiovisual (Fig. 2) sentences were presented in pseudo-random order, all interleaved in one block. In this task, participants were free to focus on one modality, or to divide attention across both modalities.

**Figure 1.**
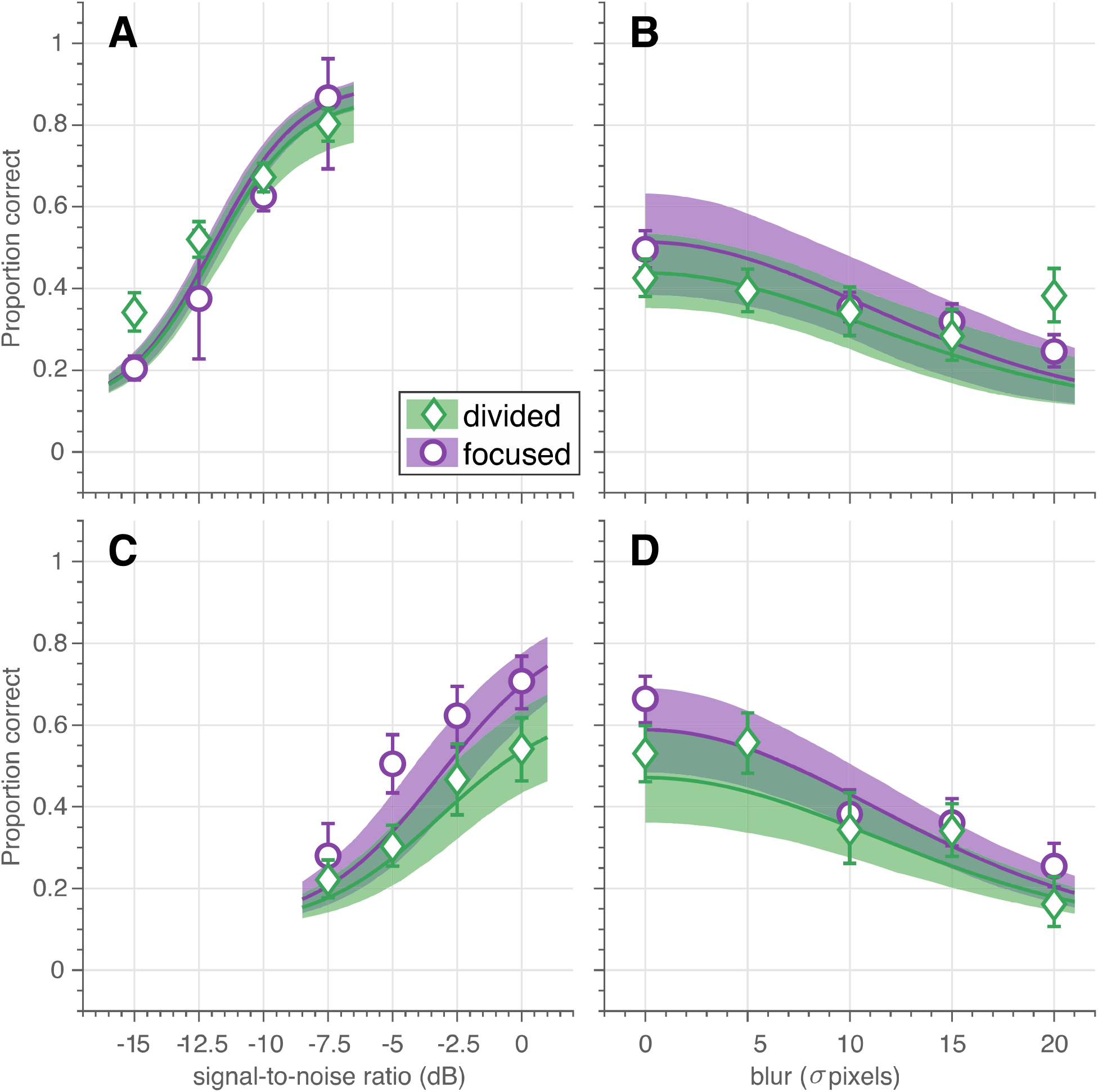
Unisensory speech recognition. (A,C) Auditory-only speech recognition (proportion correct) as a function of signal-to-noise ratio (dB) for (A) normal-hearing participants (n=14) and (C) CI users (n=7) in the focused-(purple circles) and divided-attention (green diamonds) tasks. Note that although the unisensory stimuli were the same for both tasks, CI users recognized more auditory words correctly in the focused-attention task (purple) than in the divided-attention task (green). This effect was absent for the normal-hearing participants. (B,D) Visual-only speech recognition as a function of spatial blur (in units of pixel standard deviations) for (B) normal-hearing participants and (D) CI users in the focused-(purple circles) and divided-attention (green diamonds) tasks. Note that due to the large similarity in visual recognition scores for both tasks, a psychometric curve was fitted through the combined data (black curve and patch). Symbols and bars indicate mean and 95%-confidence intervals, respectively, of the raw data (proportion correct) pooled across participants. Curves and patches indicate means and 95%-HDI, respectively, of the psychophysical-function group-level fits.

**Figure 2.**
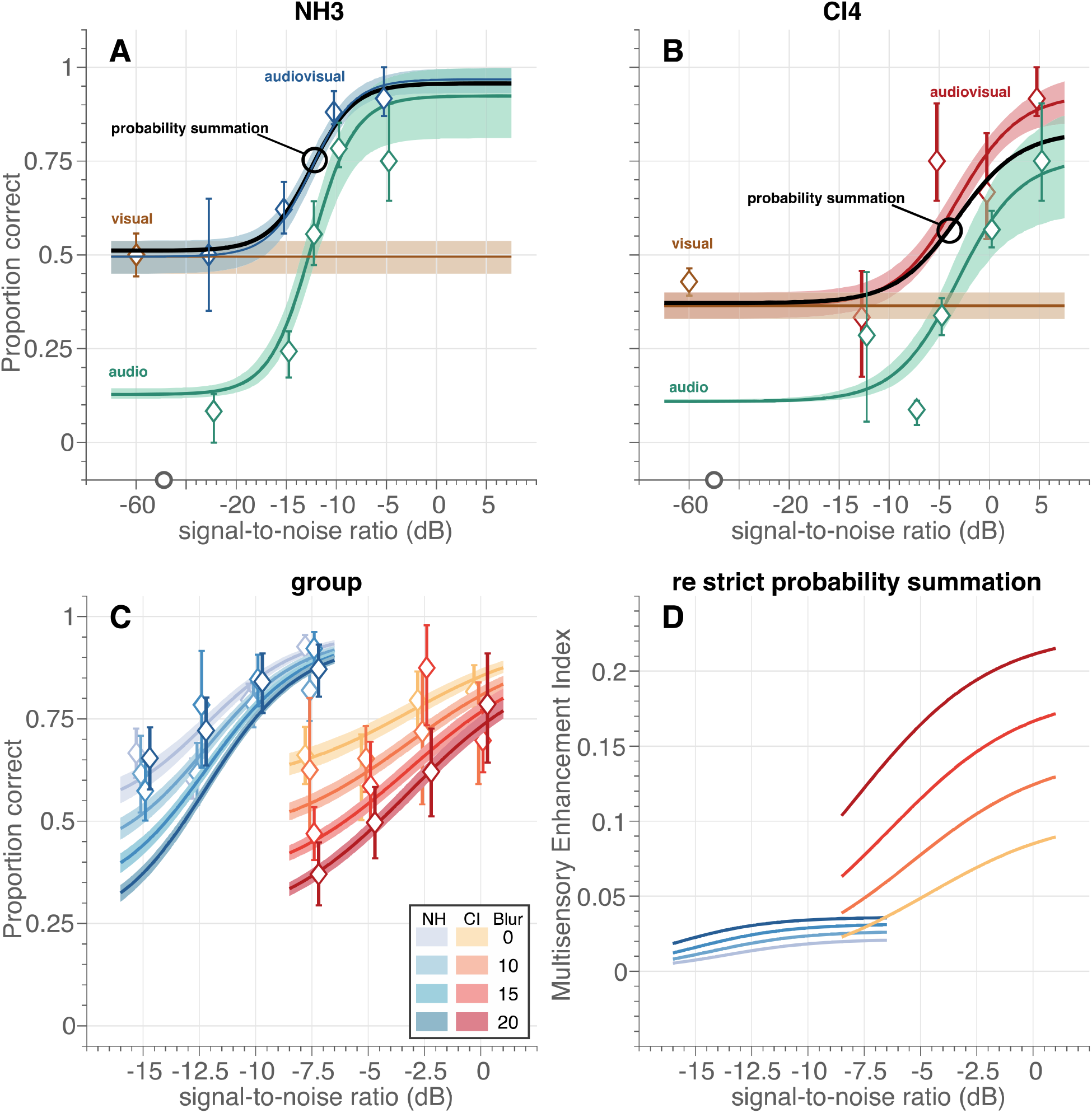
Multisensory speech recognition. Individual data and fit for (A) normal-hearing (NH) participant NH3 and (B) CI user CI4. (C) Audiovisual speech recognition scores as a function of acoustic signal-to-noise ratio (dB) for normal-hearing participants (blueish diamonds) and CI users (reddish diamonds) for four blur values (as indicated by contrast). Symbols and bars indicate mean and 95\%-confidence intervals, respectively, of the raw data (proportion correct) pooled across participants. The data was obtained (by definition) from the divided-attention task. Curves and patches indicate means and 95\%-HDI, respectively, of the psychophysical-function population fits. (D) Multisensory enhancement index as a function of acoustic signal-to-noise ratio (dB) for normal-hearing participants (blue colors) and CI users (red colors) for four blur values (as indicated by contrast). The multisensory enhancement index quantifies the multisensory enhancement of the trade-off model over strict probability summation.

To estimate parameters of interest, such as the signal-to-noise ratio and blur at which performance level was 50% and (attentional) lapse probabilities, we fitted psychophysical-function curves through the data (as fully explained in the Methods). We report on the mean and 95%-highest-density interval (HDI) of the fitted estimate distributions of the group-level parameters, and show both the fitted curves for each group and the data averaged across participants in the figures.

### Unisensory Speech Perception

When sentences were presented only acoustically (Fig. 1A,C), the two groups clearly differed in their ability to recognize words, as expected. Typically, the normal-hearing participants (Fig. 1A) recognized 50% of the words correctly in the unisensory hearing condition at a signal-to-noise ratio (auditory threshold, Eqn. 1) of −12 dB (HDI = [−12.4, −11.5] dB) vs. −3.1 dB for the CI users (HDI = [−4.4, −1.7] dB, Fig. 1C) for either of the tasks (green and purple). For both groups, the proportion of correctly recognized words strongly depended on the actual signal-to-noise ratio; to increase performance levels from 5%-to 95%-word recognition (psychometric curve width), the signal-to-noise ratio needed to be increased by 7.4 dB on average for the normal-hearing participants (HDI = [6.5, 8.5] dB; Fig. 1A) and slightly more for CI users by on average 10.4 dB (HDI = [8.8, 12.2 dB; Fig. 1C). As expected, both these results confirm that listening for CI users is considerably more difficult than for normal-hearing participants.

The parameter of main interest in this study is the lapse probability (Eqn. 3), i.e. the probability of not recognizing words even at the highest signal-to-noise ratio and without blur. Lapses occurred even in the focused-attention task as evidenced by the non-perfect performance at the highest signal-to-noise ratios; the average performance of normal-hearing participants and CI users saturated at around 90 and 84% correct, respectively (Fig.1A,C, purple; HDI = [85, 94] and [74, 92%). A larger lapse probability for the CI users compared to the lapse probability for the normal-hearing participants may be expected due to technical limits of the cochlear implant and the maximal comfortable loudness levels experienced by the CI users, but note that evidence for any difference was actually small (mean 5%, HDI = [−5, 17]%).

More importantly and more clearly, in the divided-attention task the CI users recognized 22% (HDI = [6.7, 38]%) fewer words than in the focused attention task (Fig. 1C, green vs purple). This difference was not clearly evident for the normal-hearing participants (mean difference 3.9%, HDI = [−4.0, 14] %; Fig. 1A, green vs purple). Evidence for group differences in auditory lapse probability during the divided-attention experiment was substantial (on average, the lapse probability for normal-hearing participants was 24% lower than for the CI users, HDI = [8, 41] %).

When sentences were presented only visually (Fig. 1B,D), the proportion of correctly recognized words depended on the amount of blur, and were largely similar for both groups; the visual threshold (i.e. the blur at 36% of the maximal lipreading performance, Eqn. 2 was on average 17.7 and 18.3 pixels for CI users (Fig. 1D) and normal-hearing (Fig. 1B) participants, respectively (HDI = [16.0,19.7] and [15.2, 21.8] pixels, respectively) for both tasks. Of course, lipreading abilities were far from perfect even without blurring.

No major difference in lipreading performance was observed for the visual lapse probability, so we pooled the data from both tasks to estimate this parameter. Normal-hearing participants (Fig. 1B) had a lapse in word recognition in 54% of the cases (HDI = [42, 65]%), while CI users (Fig. 1D) incorrectly recognized unblurred visual words in 46% of the cases (HDI = [36, 56]%). While one may expect CI users to be better lip-readers than normal-hearing participants, differences between groups were actually small (on average 8%, HDI = [−8, 23]%).

In summary, largely in contrast to the normal-hearing participants, the CI users experienced more speech-recognition problems when attention had to be divided between more than one sensory modality. These problems were especially conspicuous for listening, the sensory modality that faced the largest difficulties for the CI users.

### Multisensory Integration

We next analyzed whether speech perception of audiovisual stimuli would be enhanced for both groups of participants in the divided-attention task (Fig. 2). Figs. 2A and B show examples of individual participants (NH3 and CI4) in the divided-attention task at a visual blur of 10 pixels. The unisensory data and fits for these two participants (Figs. 2A,B brown and green for speech reading and listening, respectively) are in line with the group-level data and fits as described in the previous section (cf. Fig. 1, green). The audiovisual speech recognition (Fig. 2A, blue and Fig. 2B red for NH3 and CI4, respectively) outperforms or equals either unimodal speech recognition; for very low and high signal-to-noise ratios, audiovisual performance tends to equal visual or auditory performance. For intermediate signal-to-noise ratios, audiovisual performance is clearly enhanced. Such an enhancement of multisensory performance could potentially be due to mere statistical facilitation, if the participants would recognize a word by using either the available auditory, or visual information, without actually integrating both inputs. The percept is then determined by whichever sensory channel wins the race (probability summation)^9,13,20^. The audiovisual enhancement would then be fully determined by the unisensory auditory and visual recognition performance during the divided-attention task. To check for this possibility, we compared the data to the prediction from this probability-summation model (Fig. 2A,B, black curve, see Methods). For the normal-hearing participant (Fig. 2A; cf. black markers and blue curve), the model’s prediction corresponded quite well to the data. Hence, despite the improvement in audiovisual recognition rates, the normal-hearing participant did not seem to benefit from multisensory integration. In contrast, although the CI user evidently had difficulty to recognize a pure auditory speech signal in the multisensory divided-attention task (Fig. 2B, green; note the increased threshold and the larger lapse probability), they outperformed the probability-summation model for the combined audiovisual speech signals by about 10% at the highest signal-to-noise ratios (Fig. 2B, compare red vs black curves).

We quantified the audiovisual performance for all participants of both groups (visualized as a function of the acoustic signal-to-noise ratio for four different magnitudes of visual blur, Fig. 2C) by fitting a probability-summation model that was fully determined by the unisensory auditory and visual recognition performance (Eqns. 1–4). Typically, the observed multisensory enhancement should be compared to probability-summation of unisensory performance obtained from the same experimental regime, which in the current experiment would be from the divided-attention task. We term this model the strict probability-summation model. In Fig. 2C, we show the results of an alternative model, which we designate the trade-off model, that actually captures the multisensory enhancement by using the unisensory data obtained during the focused-attention task. We did this because the increased lapse probability for listening by the CI users in the divided-attention task (Fig. 1C) appeared to equal the multisensory enhancement over the strict probability-summation model (e.g. Fig. 2B, compare the red fit curve to the black curve). In essence, the difference in recognition scores between the two tasks was captured by the difference in auditory lapse probability, the single trade-off model parameter free to vary between tasks.

Nevertheless, the trade-off model describes the data for both tasks quite well (Table 1, see Methods, and Figs. 2A,B). Note that the pooled data generally appear to be at higher performance levels than the group-level fits of the trade-off model, at least for the normal-hearing participants (Fig. 2C, blue). This follows from the fact that we individualized the stimulus parameters for each participant; the data was obtained at lower signal-to-noise ratios and higher blurs more often for the better performers. The group-level fits better describe the expected overall group performance through extrapolation to a larger range of signal-to-noise ratios and blurs. By comparing the fits to the audiovisual data (Fig. 2C) to the unisensory fits (cf. Fig. 1), one can observe that audiovisual speech recognition is better than unisensory speech recognition; even at a blur of 20 pixels and a signal-to-noise ratio of −15 dB for the normal-hearing and of −7.5 dB for the CI users (around 0.2 vs 0.35 for unisensory and multisensory stimulation, respectively).

**Table 1.** Model Comparison.

| $\Delta\text{BIC}$ | Normal-hearing | CI users |
| --- | --- | --- |
| Trade-off | 0 | 0 |
| Strict | 12 | 35 |
| $R^2$ (trade-off) | 0.89 | 0.78 |
| mean signed error (trade-off) | 0.00 | +0.01 |

To illustrate the benefits of multisensory stimulation more clearly, we determined the multisensory enhancement index (MEI, Eqn. 5). This index quantifies by how much multisensory performance of the trade-off model was improved over the strict probability-summation model (Fig. 2D). A multisensory enhancement index close to zero is in line with strict statistical facilitation, while positive values are evidence for audiovisual enhancement due to multisensory integration. The index shows marginal improvement for the normal-hearing group (between 0.005-0.036, depending on signal-to-noise ratio and blur, Fig. 2D), and a far more prominent benefit for CI users that was about 4-6 times larger (0.023-0.22). A larger multisensory enhancement index for lower-informative stimuli or poorer-performing individuals would be evidence for inverse effectiveness^8,12^. This effect seemed to occur for the groups and the blurs; CI users exhibited more enhancement than the normal-hearing participants (Fig. 2D, red vs blue) and the relative multisensory improvements were largest for the highest blurs (Fig. 2D, e.g. the multisensory enhancement index for the 0-pixel blur was lower than for the 20-pixel blur, especially for the CI users). In contrast, for acoustic information a direct, rather than inverse, relationship was observed: the lowest signal-to-noise ratios elicited the smallest enhancements (Fig. 2D, the MEI curves all decline for lower signal-to-noise ratios).

## Discussion

### Summary

Results show that CI users benefit from multisensory integration in a divided-attention task (Fig. 2), but that their unisensory performance under such conditions deteriorates when compared to listening under focused attention (Fig. 1). Interestingly, their multisensory benefit matches the prediction obtained from probability summation of their (better) focused-attention performance (Fig. 2). In contrast, the normal-hearing participants do not have poorer unisensory performance in a divided-attention task, and their multisensory scores are accounted for by strict probability summation. Normal-hearing participants reached higher auditory recognition scores than the CI users. As expected, these results confirm the well-known fact that listening for CI users is considerably more difficult. Factors that likely contribute to the difficulties in understanding auditory speech in noise environments are the lack of access to finely-detailed spectral information and a limited dynamic range^21^. In contrast, CI users and normal-hearing participants had similar lipreading skills (Fig. 1B,D). This was slightly unexpected, as others have reported better lipreading abilities by CI users^18,22^. The current experiment, however, entailed recognition of a limited closed-set matrix of only 50 words. This potentially makes lipreading for normal-hearing individuals, who might be unaccustomed to lipreading in general, easier than in open sets with many more alternatives. Also, both the CI users and normal-hearing participants do have normal vision. As such, one might perhaps expect similar visual, lipreading skills.

### Attentional lapse in unisensory performance

CI users missed fewer words when they could focus on listening alone (in the focused-attention task, Fig. 1C) than in situations with uncertainty about the modality of the upcoming stimulus (in the divided-attention task). Note that this is precisely the sensory condition of every-day life. This may suggest that due to impoverished sensory information more effort is required by CI users to be able recognize speech at higher performance levels. However, the extra effort cannot be maintained by CI users if attention has to be spread out across multiple, potentially-informative sensory modalities. The CI users seem to have reached the limits of attentional resources in the divided-attention task. These limits are not reached when sensory information is not impoverished, i.e. for normal-hearing individuals and for lipreading 1A, B, D; lapse probabilities are similar across tasks).

### Multisensory integration

Following this line of reasoning, one may wonder why CI users attempt to lipread at all. Barring any other benefits, the optimal decision would be to focus on the most-informative sensory modality, and ignoring the other. Even for CI users, listening is generally (i.e. in quiet environments) the far better modality for the purposes of speech recognition. Probabilistic, uninformed switching between listening and lipreading would lead to an overall worse performance^23^. One benefit to offset this drawback could be that switching enables individuals to scan the specific environment and determine whether listening or lipreading would be the most informative modality for the given situation^24,25^. Obviously from the current experiments, another benefit could be that the detriment in listening is accompanied by an enhancement of speech recognition for multisensory stimuli. Indeed, although CI users had poorer unisensory recognition scores in the divided-attention task than in the focused attention task (Fig. 1), they outperformed the strict probability-summation model (Fig. 2D). Conversely, the normal-hearing individuals do follow strict probability summation^20^. Because of this, CI users appear to be better multisensory integrators than the normal-hearing individuals^18^ (Fig. 2D).

### Integration-attention trade-off

Intriguingly, the trade-off model suggests that the exact compensation of the listening decline (Fig. 1C) by multisensory enhancement (Fig. 2D) may be explained by an integration-attention trade-off mechanism for CI users. To benefit from multisensory integration, attention needs to be divided across all relevant signals. Only then will integration be able to enhance source identification and selection by filtering out irrelevant noise sources. The cost of this benefit is the decline in attentional amplification of unisensory signals. In our model, this is fully and solely captured by the change in auditory lapse probability (Eqn. 3), which amounted to be about 22% on average for CI users. The multisensory enhancement follows directly from this increase in lapses (through the trade-off probability-summation model, Eqns. 4 and 5); the multisensory enhancement should equal this in magnitude for the weakest visual signals and strongest auditory signals (note that the multisensory enhancement index is 0.22 for the highest blur at a signal-to-noise ratio of 0 dB), and be less for stronger visual signals and weaker acoustic signals (Fig. 2D).

## Conclusion

Normal-hearing participants can attend extensively on auditory and visual cues, while CI users need to divide their limited attentional resources across modalities to improve multisensory speech recognition - even though this leads to a degradation in unisensory speech recognition. We argue that in order to determine the acoustic benefit of a CI towards speech comprehension per se, situational factors need to be discounted by presenting speech in realistic audiovisual environments.

## Methods

### Participants

Fourteen native Dutch-speaking, normal-hearing participants (mean age: 22.3 years ± 1.8, 10 female) and 7 native Dutch-speaking, post-lingually deaf unilaterally implanted CI users (mean age 64.1 years ± 5.3, 3 female) were recruited to participate in this study. All CI users had at least one year of experience with their CI, with a mean of 3.6 years ± 1.8. Five CI users were implanted on the left. The cause of deafness was progressive sensorineural hearing loss for all but three CI users (Ménière’s disease, sudden deafness and hereditary hearing loss). Additional contralateral hearing aids were turned off during the experiment. The unaided pure tone average (range 1-4 kHz) of the non-implanted ear ranged between 70 and >120 dB Hearing Loss. However, no CI users had any speech intelligibility for words in quiet with their non-implanted ear at levels < 90 dB Sound Pressure Level (SPL). All normal-hearing participants were screened for normal hearing (within 20 dB HL range 0.5 - 8 kHz). All participants reported normal or corrected-to-normal vision. All participants gave written informed consent before taking part in the study. The experiments were carried out in accordance with the relevant institutional and national regulations and with the World Medical Association Helsinki Declaration as revised in October 2013 (Declaration). The experiments were approved by the Ethics Committee of Arnhem-Nijmegen (project number NL24364.091.08, October 18, 2011).

### Stimuli

The audiovisual material was based on the Dutch version of the speech-in-noise matrix test developed by Houben et al^26^. In general, a matrix test uses sentences of identical grammatical structure in which all available words are taken from a closed set of alternatives. The sentences are syntactically fixed (subject, verb, numeral, adjective, object), but semantically unpredictable.

The audiovisual material (Fig. 3) including the masking speech noise are reported previously^20^. Briefly, the stimulus material consisted of digital video recordings of a female speaker reading aloud the sentences in Dutch. Auditory speech (Fig. 3A,C) was presented with varying levels of acoustic background noise (Fig. 3B). Visual speech consisted of the video fragments of the female speaker (Fig. 3D). Saliency of the visual speech was altered through blurring, by filtering every image of the video with a 2-D Gaussian smoothing kernel at several pixel standard deviations.

**Figure 3.**
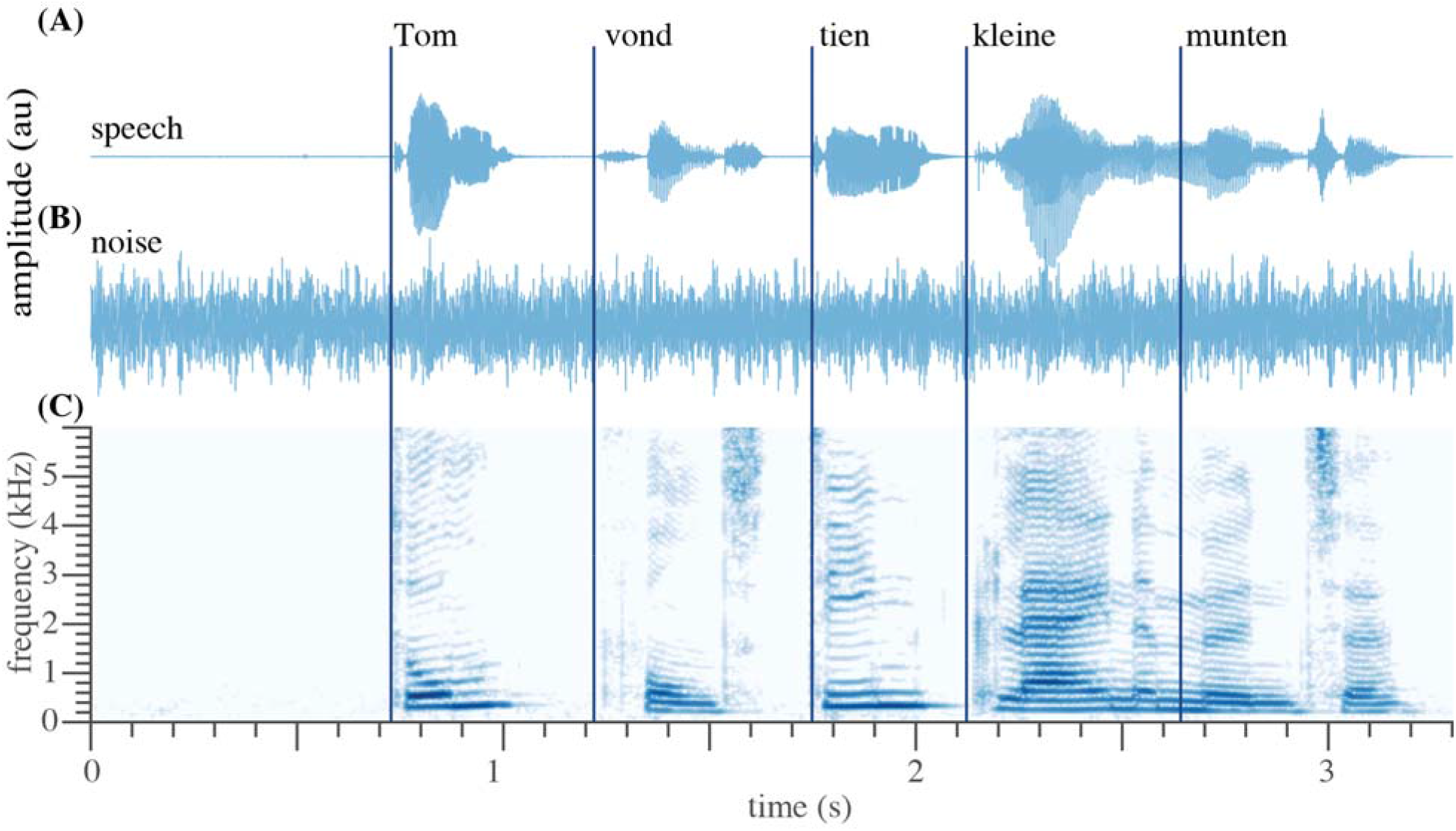
Example sentence. (A) Temporal waveform of the auditory speech signal “Tom vond tien kleine munten” (translation: Tom found ten little coins.) (B) Waveform of the auditory noise. (C) Spectrogram of the recorded sentence. (D) Five video frames around the onset of the word, untouched (top), moderately blurred (middle, 20 pixels), and extensively blurred (bottom, 70 pixels, used as a unisensory auditory condition in the divided-attention task). Dark blue lines denote the approximate onset of each individual word. Written informed consent for the publication of this image was obtained from the individual shown. Note that images in D) are not shown due to Biorxiv policy of not including images of individuals.

### Set-up

The experiments were performed in an experimental room, in which the walls and ceiling were covered with black acoustic foam that eliminated echoes for sound frequencies >500 Hz^27^. Stimulus presentation was controlled by a Dell PC (Dell Inc., Round Rock, TX, USA) running Matlab version 2014b (The Mathworks, Natick, MA, USA). were seated in a chair 1 m in front of a PC screen (Dell LCD monitor, model: E2314Hf). Sounds were played through an external PC sound card (Babyface, RME, Germany) and presented through one speaker (Tannoy, model Reveal 502) placed above the PC screen, 1 m in front of the participant (30° above the interaural plane). Speaker level was measured with an ISO-TECH Sound Level Meter, type SLM 1352P at the position of the participant’s head, using the masking noise.

### Paradigm

All participants were tested on a closed-set recognition of six Matrix lists of 20 sentences each (180 words). Participants were instructed to select words from the Matrix list which they recognized.

### Familiarization

To familiarize participants with the Matrix test procedure and to obtain an initial estimate for the auditory threshold, 40 unique auditory-only sentences were presented. The signal-to-noise ratio varied adaptively in accordance with the Brand and Kollmeier procedure^28^, and the auditory 50% speech recognition threshold was calculated as the average signal-to-noise ratio of the last nine sentences. This threshold was used to individualize the signal-to-noise ratios in focused-attention experiment. For normal-hearing participants, the speech level was fixed at 60 dB SPL, while for the CI users the noise level was fixed at 60 dB SPL. This was also true for both experiments.

### Focused-attention task: unisensory speech listening or reading

In this experiment participants listened to auditory-only sentences in one block and viewed visual-only sentences in another block. The participants were asked to accurately indicate the words (10-alternative open-ended choice per word) after each sentence. Each trial was self-paced. Participants either heard 40 or 60 unique sentences in each block.

In the auditory-only block, the auditory speech was presented in acoustic background noise with uninformative visual input (i.e. a black screen for 6 normal-hearing participants; or a heavily blurred video (70 pixel blur) for 8 normal-hearing participants and all CI users). For each sentence, the signal-to-noise ratio was pseudo-randomly picked from 4 to 12 values, that were selected individually based on the results from the adaptive tracking procedure.

In the visual-only block, the video fragments of the female speaker were shown on the screen together with the acoustic background noise and without auditory speech signal. For each sentence, the standard deviation of the Gaussian blurring kernel of the video images was pseudo-randomly picked from 5 to 10 values; the 5 most common values were 0, 6, 12, 16, and 20 pixels both for normal-hearing participants and CI users.

To avoid priming effects of sentence content (but not word content), a sentence was never repeated within a block. For each participant a different set of random signal-to-noise ratios, spatial blurs, and sentence permutations were selected. Importantly in this experiment, participants should focus on one sensory modality, and ignore the other, in order to reach maximum performance.

### Divided-attention task: multisensory speech listening and reading

In this experiment, audiovisual sentences (80 to 120 trials) were presented in one block. This experiment was conducted on another day than the focused-attention experiment. For each sentence, a visual blur and an auditory signal-to-noise ratio were chosen in pseudo-random order from five values, yielding 25 audiovisual stimulus combinations, selected in pseudo-random order. These values were selected individually based on the performance in the focused-attention experiment. We aimed for a unisensory speech-recognition performance of 0, 25, 50 and 75% for each participant, but as the maximum performance did not always reach 75%, other values were then chosen by the experimenter. The most common values were the same as for the previous experiment. In the unisensory trials of this task, the visual blur was extreme with a standard deviation of 70 pixels for the acoustic-only trials, and the auditory signal-to-noise ratio was −60 dB for the visual-only trials. Importantly, in contrast to the focused-attention task, participants could use information from both the auditory and visual modality in order to recognize words throughout most of the experiment, although some sentences were only informative in one sensory modality, but not in the other due to either extreme visual blurring (70-pixel blur) or an extremely poor acoustic signal-to-noise ratio (−60 dB signal-to-noise ratio).

### Data analysis

Proportion correct For graphical purposes, the proportion of words correct responses are plotted in raw form pooled across participants for each group as mean and 95%-HDI in Figs. 1 and 2 for the most common signal-to-noise ratios and blurs.

### Unisensory psychometric functions

To relate each participant’s responses to the intensity of the unisensory stimuli (i.e. auditory signal-to-noise ratio or visual blur), *x*, we fitted a psychometric function *F*(*x,θ*) to the unisensory data, the shape of which depended on the sensory modality, *m*. For the auditory-only data, a logistic function was fitted^20,29^:

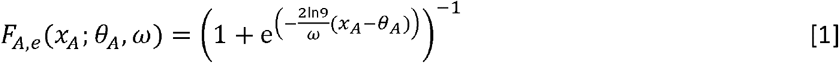

where *F_A_*(*x_A_*;*θ_A_*;*ω_A_*) characterizes the change in auditory word recognition rate as a function of the auditory signal-to-noise ratio, *x_A_*; *θ_A_* is the auditory recognition threshold for which holds *F_A_^−1^*(0.5) and *ω_A_* is the auditory recognition width, the stimulus-level range in which *F_A_* ranges from 0.1 to 0.9.

For the visual-only data, an exponential function *F_V_* was taken with only a single parameter:

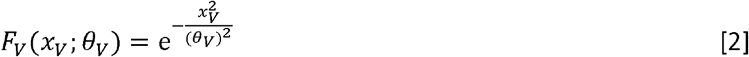

where *F_V_*(*x_V_*;*θ_V_*) characterizes the change in visual word recognition rate as a function of the visual blur, *x_V_*;*θ_V_* is the visual recognition threshold for which holds 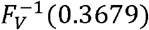, i.e. for *x_v_* = *θ_v_*. Both functions (Eqns. 1 and 2) have a sigmoidal shape and fitted the data well (i.e. Fig 1).

### Lapse

To infer the probability of correct-word recognition *Ψ*, we included a lapse probability, *λ*, to the psychometric function *F* for both modalities *m*:

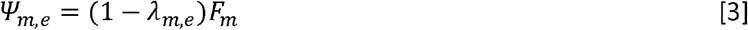

The lapse probability, *λ*, accounted for the less-than-perfect recognition probability for visual words without blurring and for auditory words at the highest signal-to-noise ratios, both for the CI users and the normal-hearing participants. With probability *λ*_(*m,e*)_ a participant has a momentary lapse (i.e. makes a choice independent of stimulus intensity) for modality *m* during experiment *e*. With probability (1 − *λ*) the participant does not have a lapse and has a chance of *F_m_* to give the correct answer. The lapse probability could reflect several issues: e.g. a momentary lapse of attention, blinking during the visual trials, or the lack of increase in information with increasing stimulus intensity due to for example processing issues of the cochlear implant.

Crucially, the estimate for the lapse probability was, at first, inferred separately for the experimental tasks (focused-attention vs divided), as we hypothesized that the separate tasks could differentially affect attentional demands, potentially leading to observed differences in attentional lapses.

We modified this slightly, as we observed no significant differences in the visual lapse probability between experimental tasks (Fig. 1). Thus, the final fitted model (Eqns. 1–3), as reported here, included the auditory lapse probability as the only parameter that was free to vary between experimental tasks. Constraining the model in such a way had no effect on the conclusions.

### Multisensory psychometric function defined by probability summation

We modelled the audiovisual speech recognition as a mere statistical-summation effect that is distinct from true neural audiovisual integration. In this model of probability summation (see Introduction), participants recognize a word from either the auditory-only or the visual-only condition, which are considered independent processing channels. Thus, if a subject fails to recognize a word from either one of the modalities, the probability of failure is (1−*Ψ_A_*) × (1−*Ψ_V_*). It then follows that the probability of word recognition in the presence of the two modalities without integration is given by:

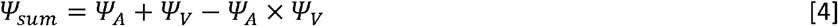

where *Ψ_sum_* is the probability to successfully recognize a word according to the summation model, *Ψ_A_* is the probability to recognize an auditory word in the auditory-only condition, and *Ψ_V_* is the probability of recognizing a visual word. From this, one can observe that having both modalities available, rather than one, automatically increases the probability of stimulus recognition.

We chose to fit this model because previous evidence^20^ showed that speech recognition of the audiovisual materials could be described well by probability summation. Importantly, the data was accurately fitted by this model (see also the section on Model Selection), with one caveat: the fit was better if the lapse probabilities for the audiovisual stimuli (by definition, only presented in the divided-attention task) was set to equal the lapse probabilities as found in the focused-attention task.

This meant that model could only predict an enhancement of speech recognition for multisensory stimuli through a combination of mere statistical facilitation and a change in auditory lapse probability across experimental tasks. To visualize this (Fig. 2D), we determined the multisensory enhancement index, MEI:

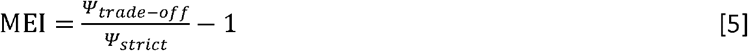

with *Ψ_strict_* and *Ψ_trade-off_* the probability to successfully recognize a word according to the summation model with an auditory lapse probability taken from the divided-attention (strict) and focused-attention (trade-off) tasks, respectively. An MEI close to zero is in line with statistical facilitation, and no change in lapse probability. Positive values are evidence for an observed multisensory enhancement and an increased auditory lapse probability.

### Guess probability

We also included a guess rate of 10% that accounts for a fixed probability of 0.1 of correctly choosing one of the ten alternatives by chance alone (0.9*Ψ* + 0.1). This was the same for every participant, modality and experimental task, as it depended on the design of the Matrix test itself.

### Approximate Bayesian inference

Parameter estimation was performed using approximate Bayesian inference. The models described by eqns. 1–4 were fitted on all data simultaneously. The parameters were estimated for every participant, which depended on the estimation of overarching group parameters, separately for the normal-hearing participants and CI users, in a hierarchical fashion.

The estimation procedure relied on Markov Chain Monte Carlo (MCMC) techniques. The estimation algorithms were implemented in JAGS^30^ through matJAGS^31^. Three MCMC chains of 10,000 samples were generated. The first 10,000 samples were discarded as burn-in. Convergence of the chains was determined visually, by checking that the shrink factor 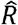 is less than 1.1 and by checking that the effective sample size is larger than 1000^32^. From these samples of the posterior distributions, we determined the mean and the 95%-HDI as a centroid and uncertainty estimate of the parameters, respectively.

### Model Selection

To test for the appropriateness of the models in eqns. 1–4, we compared them against less-restrictive models. To that end, we performed a qualitative check via visual inspection (c.f. Figs. 1 and 2), but we also quantitatively determined the Bayesian Information Criterion (BIC) for each model:

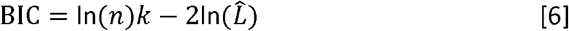

where *k* denotes the number of parameters of the model, *n* the number of samples and 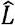 the maximized value of the binomial likelihood function.

## Author contributions

LR, JO and MW designed research; LR performed research; MW analyzed data; LR, JO and MW wrote the paper; LR and MW drafted the initial research concept.

## Acknowledgments

We thank Günther Windau, Ruurd Lof, Stijn Martens, and Chris-Jan Beerendonck for their valuable technical assistance, speech-therapist Jeanne van der Stappen for being the speaker in the audiovisual material, Eefke Lemmen for video editing, and Roos Cartignij for help in data acquisition. We are grateful to Ad Snik and Emmanuel Mylanus for providing valuable comments on earlier versions of this manuscript.

## Funding

This research was supported by EU Horizon 2020 ERC Advanced Grant ORIENT (grant 693400, JO), Cochlear Benelux NV (LR, MW), the Radboud University Medical Center (LR), and Radboud University (MW). The funders had no role in study design, data collection and analysis, decision to publish, or preparation of the manuscript.

## Author declaration

The authors declare no competing interest.

## Data deposition

The data have been deposited in the Donders Institute for Brain, Cognition and Behavior Data Repository at https://doi.org/10.34973/jy8p-dw52.

## Notes

### Competing Interest Statement

The authors have declared no competing interest.

